# Parkinson’s disease associated mutation E46K of α-synuclein triggers the formation of a novel fibril structure

**DOI:** 10.1101/870758

**Authors:** Kun Zhao, Yaowang Li, Zhenying Liu, Houfang Long, Chunyu Zhao, Feng Luo, Yunpeng Sun, Youqi Tao, Xiao-dong Su, Dan Li, Xueming Li, Cong Liu

**Affiliations:** Interdisciplinary Research Center on Biology and Chemistry, Shanghai Institute of Organic Chemistry, Chinese Academy of Sciences, Shanghai 201210, China; University of Chinese Academy of Sciences, Beijing 100049, China; Key Laboratory of Protein Sciences (Tsinghua University), Ministry of Education, School of Life Sciences, Tsinghua University, Beijing 100084, China; Bio-X Institutes, Shanghai Jiao Tong University, Shanghai 200030, China; State Key Laboratory of Protein and Plant Gene Research, and Biomedical Pioneering Innovation Center (BIOPIC), School of Life Sciences, Peking University, Beijing 100871, China

## Abstract

α-Synuclein (α-syn) amyloid fibril, as the major component of Lewy bodies and pathological entity spreading in human brain, is closely associated with Parkinson’s disease (PD) and other synucleinopathies. Several single amino-acid mutations (e.g. E46K) of α-syn have been identified causative to the early onset of familial PD. Here, we determined the cryo-EM structure of a full-length α-syn fibril formed by N-terminal acetylated E46K mutant α-syn (Ac-E46K). The fibril structure represents a new fold of α-syn, which demonstrates that the E46K mutation breaks the electrostatic interactions in the wild type (WT) α-syn fibril and thus triggers the rearrangement of the overall structure. Furthermore, we show that the Ac-E46K fibril is less resistant to harsh conditions and protease cleavage, and more prone to be fragmented with a higher capability of seeding fibril formation than that of the WT fibril. Our work provides a structural view to the severe pathology of the PD familial mutation E46K of α-syn and highlights the importance of electrostatic interactions in defining the fibril polymorphs.

## INTRODUCTION

Deposition of α-syn amyloid fibrils in Lewy bodies (LB) and Lewy neuritis (LN) is a common histological hallmark of Parkinson’s disease (PD) and synucleinopathies such as dementia with Lewy bodies (DLB) and multiple system atrophy (MSA)^1–3^. Accumulating evidence show that α-syn amyloid fibrils serve as prion-like seeds for the propagation and cell-to-cell transmission of the pathological entity of α-syn^4,5^. The spread of pathological α-syn fibril is closely associated with the disease progression^6,7^. Several single point mutations and genomic duplication or triplication of *SNCA*, the gene encoding α-syn, have been identified to be causative to the familial forms of the diseases with a broad spectrum of distinctive clinical symptoms ^8–12^. Moreover, the hereditary mutations exhibit exacerbated pathology in various cellular and PD-like animal models^13,14^. Intriguingly, the cryo-EM structure of full-length wild-type (WT) α-syn fibril demonstrates that in the five mutation sites found in familial PD, four of them, i.e. E46K, A53T/E, G51D and H50Q, are located at the protofilamental interface of the WT fibril^15,16^, which indicates that these mutations may alter the fibril structure and consequently influence α-syn amyloid aggregation and PD pathology.

α-Syn hereditary mutation E46K was originally identified from a Spanish family with autosomal dominant parkinsonism^11^. The clinical phenotype of E46K patient features rapid and severe disease progression with early onset of the parkinsonism of DLB^11^. Previous studies have shown that the E46K mutation can enhance α-syn fibril formation and α-syn pathology *in vitro* and in cultured cells^13,17,18^. Solid state NMR study has shown that the E46K mutation causes large conformational changes of the fibril structure^19^.

In this work, we determined the cryo-EM structure of the N-terminally acetylated E46K α-syn (Ac-E46K) fibril at an overall resolution of 3.37 Å. The structure reveals a distinct fold of α-syn, in which the electrostatic interactions in the WT fibril are broken and reconfigured due to the E46K mutation. We further show that the Ac-E46K fibril is less stable than the WT fibril under harsh conditions and protease cleavage, while the mutant fibril is more efficient in seeding amyloid fibril formation. This work provides a structural mechanism for the influence of E46K on the amyloid fibril formation of α-syn, and suggests that electrostatic interactions may serve as one of the driving force for the polymorphic fibril formation of α-syn.

## RESULTS

### E46K mutation alters the morphology and stability of α-syn fibril

To investigate the influence of E46K on α-syn fibril formation, we prepared recombinant full-length α-syn with E46K mutation. Since *in vivo* α-syn is generally acetylated at the N terminus^20,21^, we modified the recombinant α-syn with N-terminal acetylation (Supplementary Fig. 1). The recombinant N-acetylated E46K α-syn (termed as Ac-E46K) formed amyloid fibrils after incubation in the buffer containing 50 mM Tris, 150 mM KCl, pH 7.5, at 37°C with agitation for 7 days. We first used the atomic force microscopy (AFM) to characterize the fibril morphology. The result showed that the Ac-E46K fibril features a different fibril structure from the N-acetylated WT (Ac-WT) fibril formed under the same condition ^15^ (Supplementary Fig. 2). The Ac-E46K fibril is approximately two times more twisted than the Ac-WT fibril with a pitch of ~ 64 nm in comparison with ~ 120 nm of the Ac-WT. Moreover, the Ac-E46K fibril features a right-handed helical twist (Supplementary Fig. 2), which is distinct from the left-handed twist commonly found in the WT and other PD-familial mutant α-syn fibrils^15,16,22–24^. The different fibril morphologies suggest different properties of the Ac-E46K and Ac-WT fibrils, which may associate with their different pathologies.

To identify the different properties of the Ac-E46K and Ac-WT fibrils, we first sought to test the stability of the fibrils by cold denaturation^25^. The fibrils were cooled down and incubated at 0 °C for 48 h. Fibril disassociation was assessed by the loss of β structures monitored by circular dichroism (CD) spectroscopy. The result showed that the Ac-E46K fibril denatured significantly faster than that of the Ac-WT fibril (Fig. 1a). In addition, we found that during the experiment, the Ac-E46K fibril stored at −80 °C underwent apparent denaturation after thawing; in contrast, the Ac-WT fibril were generally stable. To quantitatively measure the stability of the fibrils upon cycles of freeze-thaw, we flash-froze the α-syn fibrils in liquid nitrogen and thawed the fibrils at room temperature in water bath. CD spectra showed that in eight cycles of freeze-thaw, the structure of Ac-WT fibril was well maintained with a consistent content of β structures (Fig. 1b). In contrast, the Ac-E46K fibril gradually lost its β structures as the freeze-thaw cycles increased, indicating the disassembly of amyloid fibrils (Fig. 1b). Thus, these results indicate that the Ac-E46K fibril is less stable than the Ac-WT fibril.

**Fig. 1.**
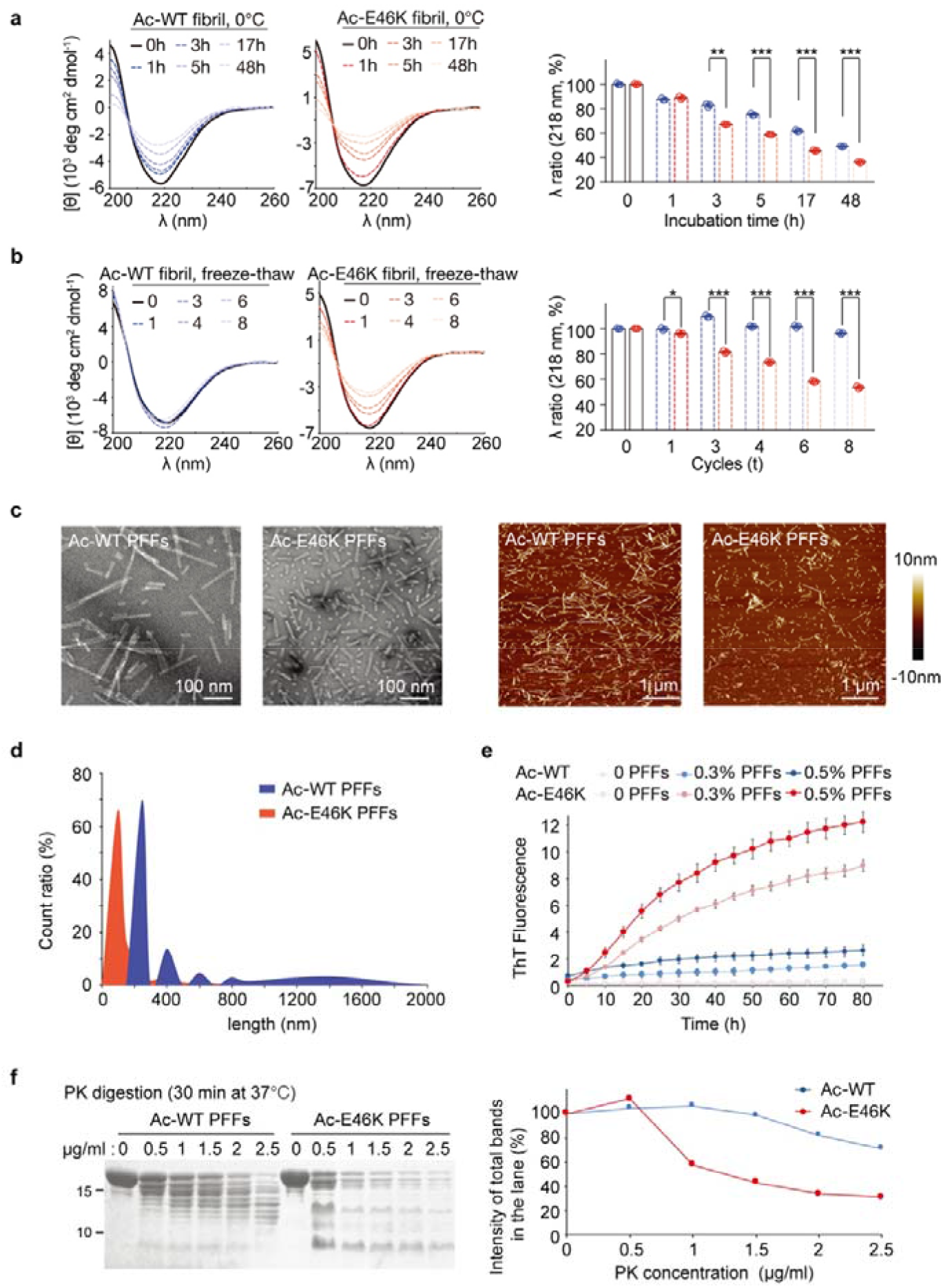
Comparison of the stability of the Ac-WT and Ac-E46K α-syn fibrils. **a** Cold denaturation of 20 μM Ac-WT (blue) and Ac-E46K (red) α-syn fibrils. CD spectra monitored at 0 □ at different incubation time are shown on the left. CD signals at 218 nm are analyzed on the right. Data are shown as mean ± S.D., with n = 3 for both panel a and b. \**p* < 0.05, \*\**p* < 0.01 and \*\*\**p* < 0.005. **b** Freeze-thaw denaturation of Ac-WT (blue) and Ac-E46K (red) fibrils. CD spectra obtained at different freeze-thaw cycles are shown on the left. CD signals at 218 nm are analyzed on the right. **c** Negative-staining TEM (left) and AFM (right) images of 5 μM Ac-WT and Ac-E46K α-syn PFFs after sonication. **d** Size distribution of sonicated Ac-WT and Ac-E46K PFFs. The lengths of the sonicated PFFs are measured by AFM. 300 fibrils were measured for each fibril sample. **e** Different seeding effects of the sonicated Ac-WT and Ac-E46K PFFs for their self-proliferation by the ThT kinetic assay. Concentrations of the PFF seeds are indicated. **f** PK digestion of the Ac-WT and Ac-E46K fibrils. The fibrils were incubated with indicated concentrations of PK at 37 □ for 30 min (left). The intensity of the total protein bands in each lane of the SDS-PAGE is analyzed on the right.

Since α-syn fibrils propagate by seeding the amyloid fibril formation in the transfected neurons, we attempted to compare the seeding property of the Ac-E46K and Ac-WT fibrils. We prepared the fibril seeds by sonication of the preformed fibrils (PFFs). Intriguingly, we noticed that under the same sonication condition, the Ac-E46K fibril broke into significantly shorter fragments than that of the Ac-WT fibril visualized by negative-staining transmission electron microscopy (TEM) and AFM (Fig. 1c, d), which is consistent with the stability tests, and more importantly may reflect an increased pathology of the Ac-E46K fibril since fibril fragmentation has been found as a key factor in nucleation-depended fibrillation^26^. Next, we used the fragmentated PFFs to seed Ac-E46K and Ac-WT, respectively, and monitored the amyloid fibril formation by the ThT fluorescence assay and TEM. The result showed that under the tested condition, there was no obvious fibril formed by the Ac-E46K or Ac-WT α-syn without seeding (Fig. 1e, Supplementary Fig. 3). While, in the presence of fibril seeds, fibril formation was observed, and the Ac-E46K fibril seeds showed markedly higher seeding efficiency than that of the Ac-WT (Fig. 1e, Supplementary Fig. 3), which suggests that the E46K fibril may have a higher ability of propagation and thus may be more pathological than the WT fibril.

Furthermore, we compared the stability of the fibrils under proteinase K (PK) digestion. The result showed that the Ac-E46K fibril is digested significantly faster than that of the Ac-WT fibril (Fig. 1f). Together, these data indicate that the Ac-E46K fibril is less stable than the Ac-WT fibril in both physical and chemical treatments and more efficient in fragmentation and propagation.

### Structure determination of the Ac-E46K α-syn fibril by cryo-EM

To understand the structural basis underlying the different properties between the Ac-E46K and Ac-WT fibrils, we sought to determine the atomic structure of Ac-E46K fibril by cryo-EM. The Ac-E46K fibril sample was fixed on the carbon grid and frozen in liquid ethane. The cryo-EM micrographs were acquired at 105,000× magnification on the 300 keV Titan Krios microscope equipped with the K2 Summit camera. 13,064 fibrils picked from 754 micrographs were used for the reconstruction of the Ac-E46K fibril (Table 1). The Ac-E46K fibril sample is morphologically homogeneous with a one dominant species in the 2D classification of the fibrils (Supplementary Fig. 4). After helical reconstruction of the dominant fibril species by Relion, we obtained a 3D density map of the Ac-E46K fibril to an overall resolution of 3.37 Å (Supplementary Fig. 5). The density map showed a right-handed helix with a width of ~10 nm and a helical pitch of ~68 nm (Fig. 2), which is consistent with the AFM measurement (Supplementary Fig. 2). The fibril contains two protofilaments that intertwine along an approximate 2-fold screw axis (Fig. 2). The helical twist between α-syn subunits is −179.37° and the helical rise is 2.38 Å (Fig. 2b, c, Table 1).

**Fig. 2.**
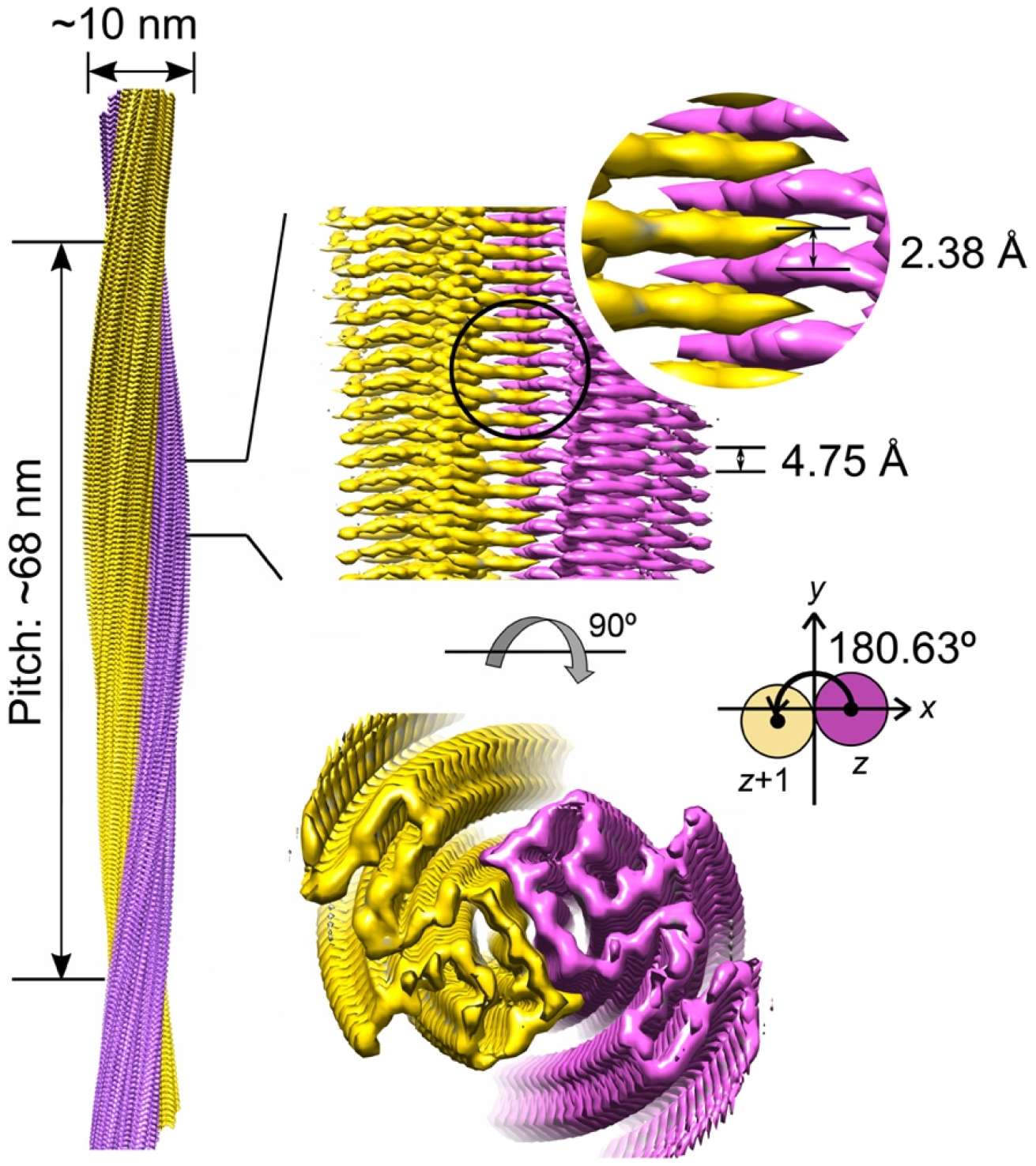
Cryo-EM 3D reconstruction density map of the Ac-E46K α-syn fibril. Fibril width, pitch length, helical rise and twist angle are indicated. The twist angle is graphically illustrated. The two intertwining protofilaments are colored in yellow and purple, respectively.

**Table 1.**
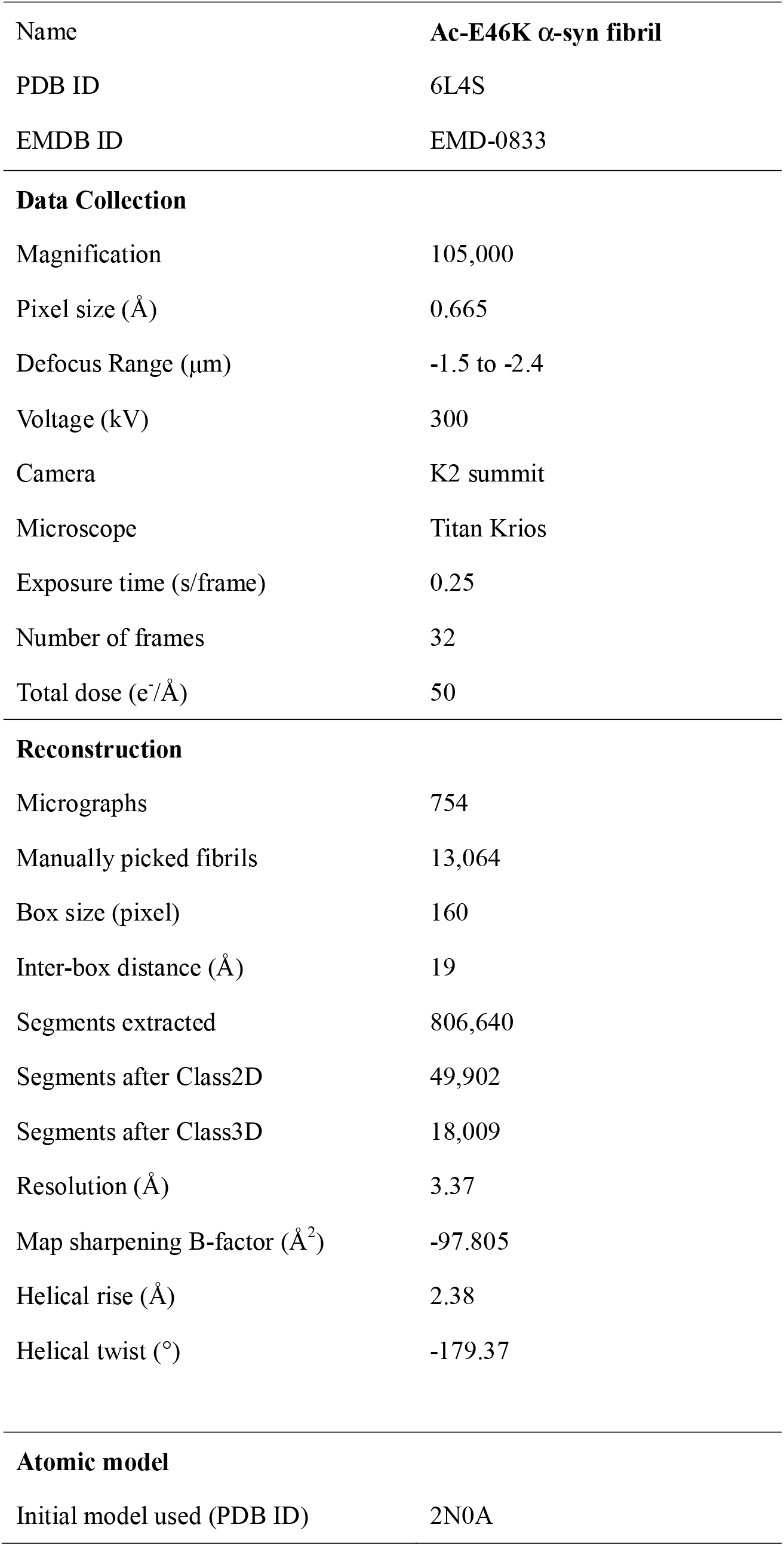
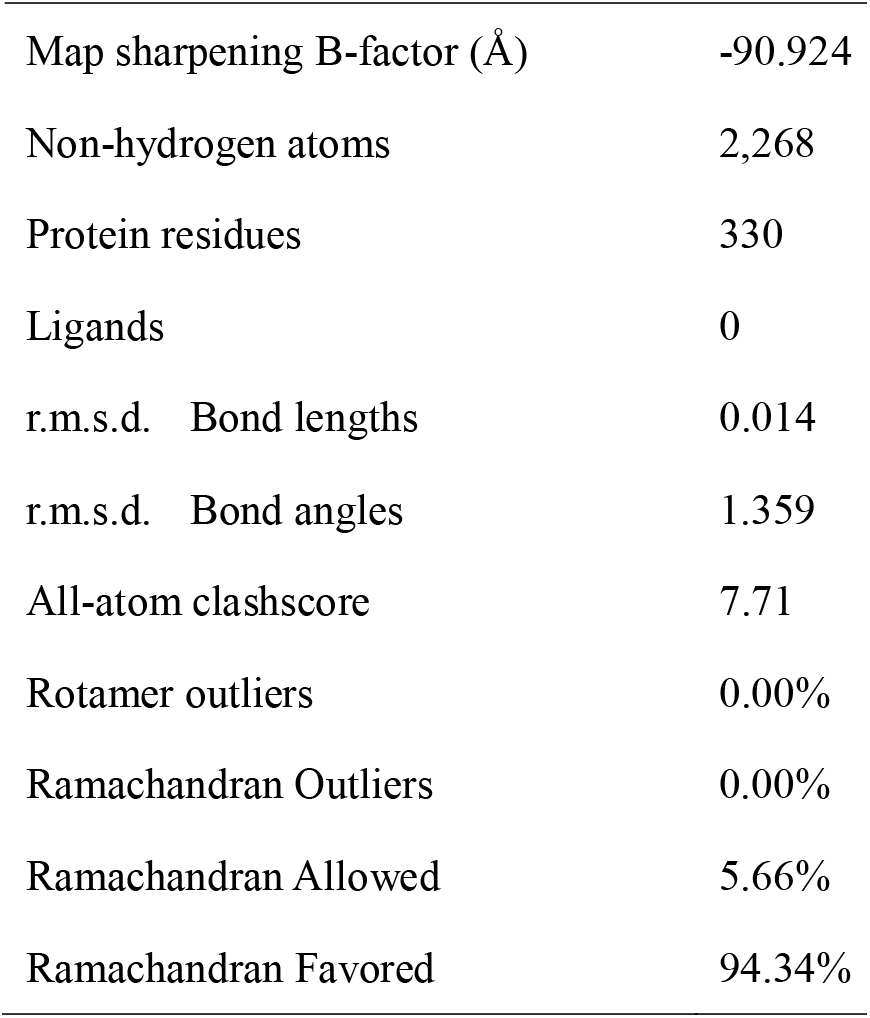
Statistics of cryo-EM data collection and refinement.

|  |  |
| --- | --- |
| Name | Ac-E46K α-syn fibril |
| PDB ID | 6L4S |
| EMDB ID | EMD-0833 |
| **Data Collection** |  |
| Magnification | 105,000 |
| Pixel size ($\text{\AA}$) | 0.665 |
| Defocus Range ($\mu\text{m}$) | -1.5 to -2.4 |
| Voltage (kV) | 300 |
| Camera | K2 summit |
| Microscope | Titan Krios |
| Exposure time (s/frame) | 0.25 |
| Number of frames | 32 |
| Total dose ($\text{e}^-/\text{\AA}$) | 50 |
| **Reconstruction** |  |
| Micrographs | 754 |
| Manually picked fibrils | 13,064 |
| Box size (pixel) | 160 |
| Inter-box distance ($\text{\AA}$) | 19 |
| Segments extracted | 806,640 |
| Segments after Class2D | 49,902 |
| Segments after Class3D | 18,009 |
| Resolution ($\text{\AA}$) | 3.37 |
| Map sharpening B-factor ($\text{\AA}^2$) | -97.805 |
| Helical rise ($\text{\AA}$) | 2.38 |
| Helical twist ($^\circ$) | -179.37 |
| **Atomic model** |  |
| Initial model used (PDB ID) | 2N0A |
| Map sharpening B-factor (Å) | -90.924 |
| Non-hydrogen atoms | 2,268 |
| Protein residues | 330 |
| Ligands | 0 |
| r.m.s.d. Bond lengths | 0.014 |
| r.m.s.d. Bond angles | 1.359 |
| All-atom clashscore | 7.71 |
| Rotamer outliers | 0.00% |
| Ramachandran Outliers | 0.00% |
| Ramachandran Allowed | 5.66% |
| Ramachandran Favored | 94.34% |

### The Ac-E46K fibril exhibits a novel fold of α-syn fibril

The high quality cryo-EM density map allowed us to unambiguously build an atomic structure model for the Ac-E46K fibril (Fig. 3a). This structure represents the fibril core (FC) of the Ac-E46K fibril consisting of residues 45-99 out of a total of 140 amino acids of α-syn (Fig. 3a), which is slightly smaller than the Ac-WT FC consisting of residues 37-99 formed under the same condition^15^. While, similar to that of the WT fibril, the N-and C-termini of Ac-E46K α-syn remain flexible and are not visible by cryo-EM.

**Fig. 3.**
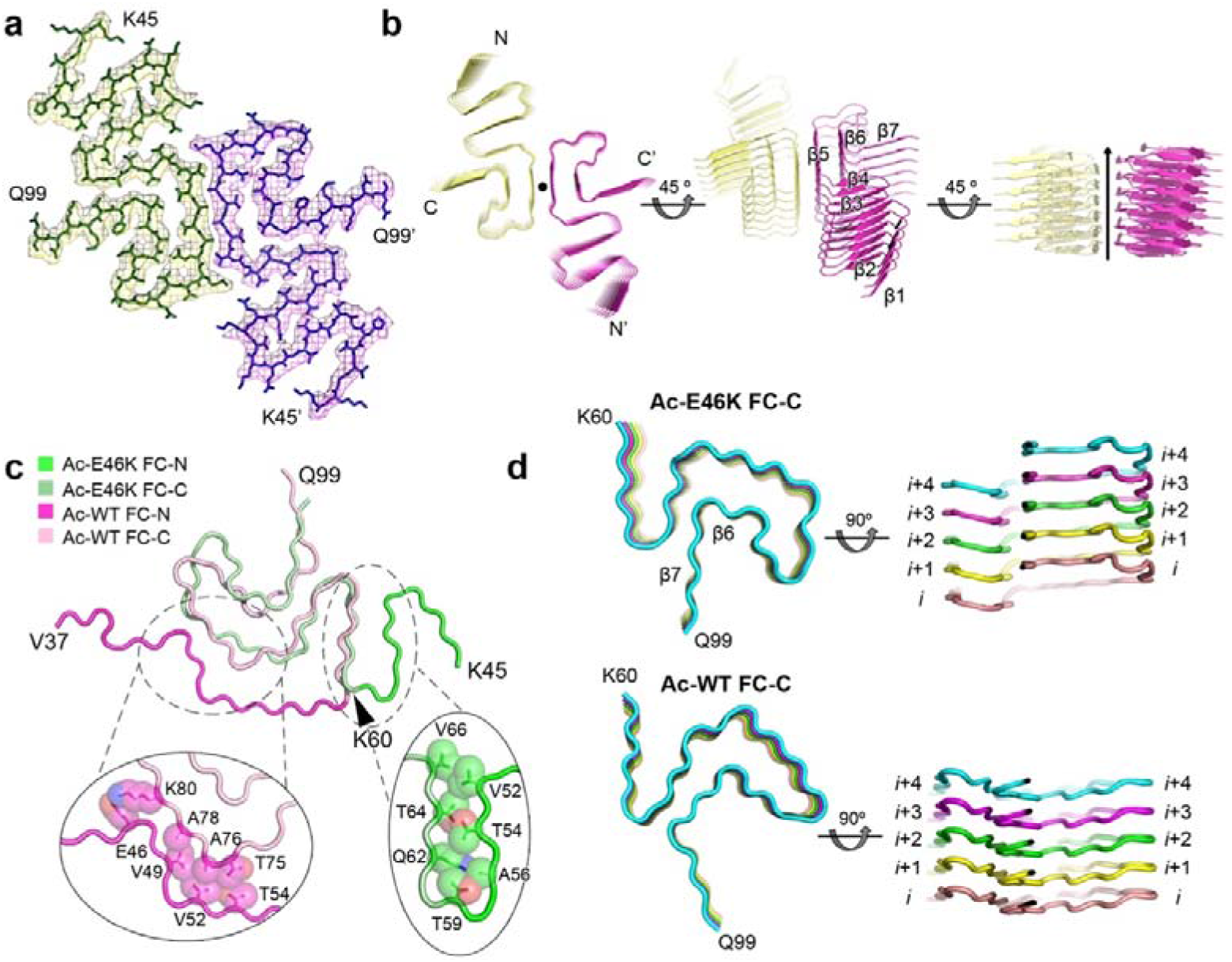
Cryo-EM structure of the Ac-E46K α-syn fibril. **a** Top view of the Ac-E46K fibril. One layer of the structure is shown, which consists of two α-syn molecules covering residues 45-99. The two molecules are colored differently. **b** Views of six layers of the Ac-E46K fibril are shown in cartoon. The two protofilaments are colored differently. The fibril axis is indicated. The β strands are numbered and labeled. **c** Overlay of the α-syn subunit structures of the Ac-E46K and Ac-WT fibrils. The two structures diverge mainly at the FC-N region. The interfaces between the FC-N and FC-C regions are shown in the zoom-in views. The residues involved in the interfaces are labeled and their side chains are shown in spheres. **d** Comparison of the FC-C region of the Ac-E46K and Ac-WT fibrils. Five layers of a single protofilament is shown. The structures are colored by different layers. The FC-C region adopts similar topology in the two structures, while their structural arrangements in the two fibrils are markedly different. The FC-C of the Ac-WT fibril folds in a flat layer. In contrast, in the Ac-E46K fibril, β6 and β7 swap to the next α-syn molecule in the neighboring layer.

The Ac-E46K FC exhibits a serpentine fold, in which the topology of the β strands is different from any reported α-syn fibril structure^15,16,22–24,27,28^ (Fig. 3b). Compared to the structure of Ac-WT α-syn fibril, the major difference lies in the N-terminal region of the FC (termed as FC-N), which covers residues 37-59 in the Ac-WT fibril or 45-59 in the Ac-E46K fibril. In the Ac-WT structure, the FC-N region stretches and packs around the rest of the α-syn molecule (termed as FC-C) via interactions with T75, A76, A78 and K80 of FC-C (Fig. 3c). In contrast, in the Ac-E46K structure, this segment forms a β hairpin and extends from the FC-C with interactions to the side chains of Q62, T64 and V66 (Fig. 3c).

Aside from the large conformational change of the FC-N, the FC-C regions of the Ac-E46K and Ac-WT fibrils adopt a similar topology with a Greek key-like fold (Fig. 3c); however, the two FC-C structures are very different with an r.m.s.d. (Cα atoms) of 5.558 Å (Supplementary Fig. 6). In the Ac-WT fibril, α-syn subunits fold in a flat layer; in contrast, in the Ac-E46K fibril, β6 and β7 of the FC-C swap to the next layer to form inter-molecular side-chain interactions within the same protofilament (Fig. 3d). Thus, the single mutation of E46K entirely changes the α-syn fibril structure.

### E46K mutation rearranges the electrostatic interactions in the α-syn fibril

Since the overall structural rearrangement of α-syn fibril is initiated by one single mutation of E46K, we looked close to residue 46 in the Ac-WT and Ac-E46K fibrils. In the Ac-WT fibril, E46 forms a salt bridge with K80, which together with the salt bridge formed by E61 and K58, locks the FC-N with FC-C (Fig. 4a, b). In contrast, in the Ac-E46K fibril, the E46K mutation breaks the salt bridge of E46-K80, which results in the breaking of the other two essential electrostatic interactions (K58-E61 and K45-H50-E57) (Fig. 4a, b). Alternatively, in the Ac-E46K fibril, K80 forms inter-molecular salt bridge with E61 of the opposing α-syn subunit to involve in the fibril interface, and K45 directly forms salt bridge with E57 to stabilize the β-hairpin conformation of the FC-N (Fig. 4a, b).

**Fig. 4.**
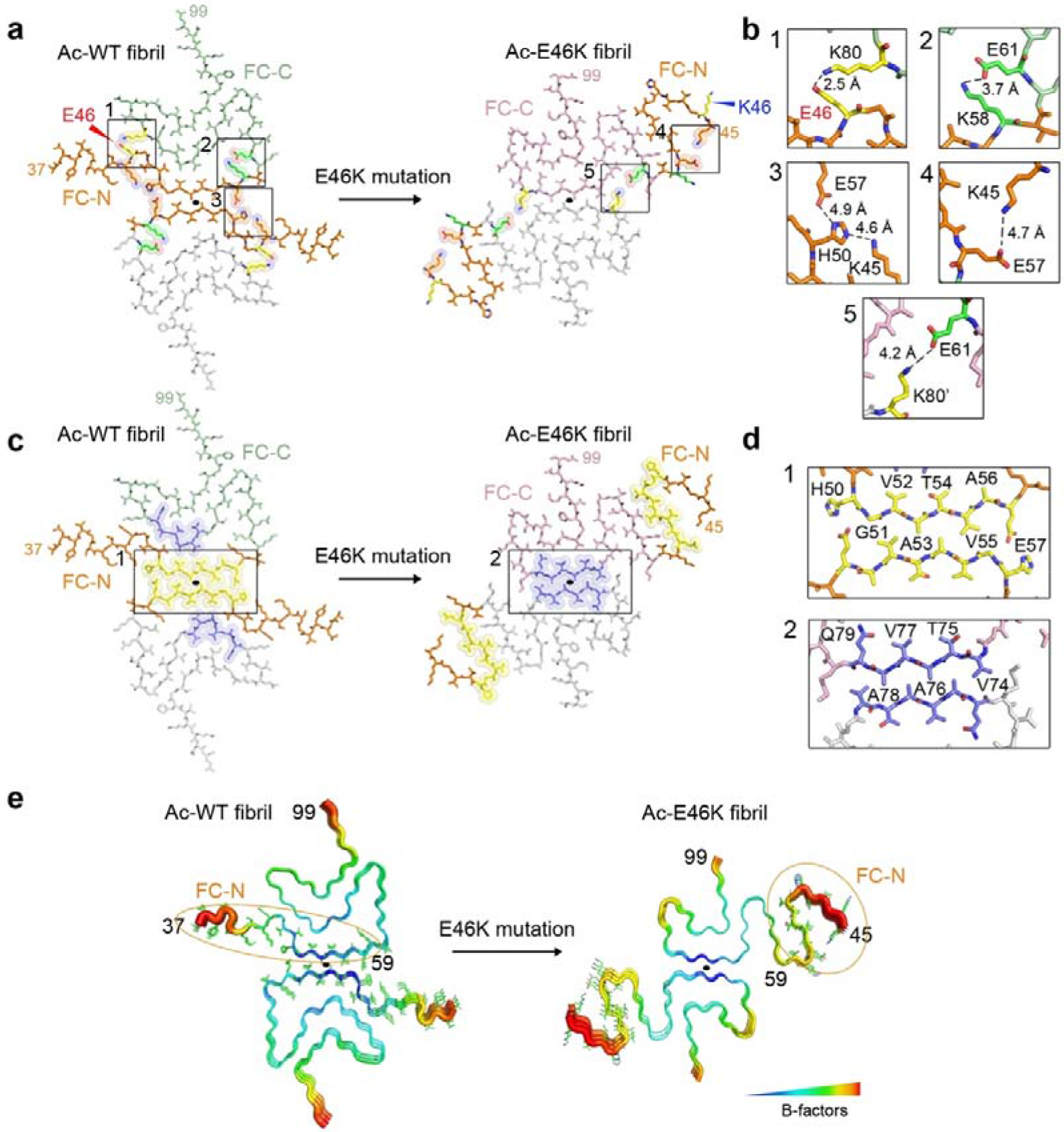
Rearrangement of the α-syn fibril structure triggered by E46K mutation. **a** Electrostatic interactions in the Ac-WT and Ac-E46K fibrils. K/E pairs that form salt bridges are highlighted with spheres and shown in zoom-in views in (b). The FC-N region (residues 37/45-59) is colored in orange. The mutation site E46/K46 is highlighted in red/blue. The black solid ellipses in all panels indicate the symmetric axis of the dimeric α-syn structure. **b** Zoom-in views of the corresponding regions in (a). Distances of the electrostatic interactions are indicated, and involved residues are labeled. The mutation site E46 is highlighted in red. **c** Protofilamental interfaces of the Ac-WT and Ac-E46K fibrils. Residues involved in the interfaces are highlighted with spheres. The fibril interface of Ac-WT fibril is colored in yellow; that of the Ac-E46K fibril is in blue. The interfaces are zoomed in (d). **d** Zoom-in views of the fibril interfaces. Interface residues are labeled. **e** Conformational change of the FC-N region results in a decreased stability of this region. Three layers of the α-syn fibrils are shown in B-factor putty. The FC-N region is shown in lines and highlighted with orange ellipses.

The E46K mutation also leads to the disruption of the WT protofilament interface (Fig. 4c, d). Upon the flipping away of the FC-N from the fibril interface, a new fibril interface is formed. The exposed segments 74-79 of opposing α-syn subunits form typical class I parallel in-register homo-steric zippers to mediate the inter-protofilamental interactions (Fig. 4d). The steric-zipper interface is stabilized by the hydrophobic interactions between the side chains of V74, A76 and A78 and flanked by the inter-molecular salt bridges formed by K80-E61’, which further lock the two protofilaments (Fig. 4a).

The release of FC-N from the fibril interface in the Ac-E46K fibril increases the flexibility of this segment, which exhibits higher B-factors and poor density for tracing the N-terminal residues 37-44 that are visible in the Ac-WT FC (Fig. 4c). Thus, the relatively loose packing of the Ac-E46K fibril in comparison with that of the Ac-WT fibril may explain the decreased stability and increased sensitivity of the Ac-E46K fibril to the environment, which may associate with the pathology of this PD familial mutation.

## DISCUSSION

Structural polymorphism is a common characteristic of amyloid fibrils formed by different amyloid proteins such as α-syn, tau and Aβ^29–31^. Different fibril polymorphs may represent different pathological entities that are associated with different subsets of neurodegenerative diseases^32–34^. Determination of polymorphic amyloid fibril structures is useful for the mechanistic understanding of amyloid pathology. In this work, we report the cryo-EM structure of an α-syn fibril with N-terminal acetylation and familial mutation E46K found in parkinsonisms. The Ac-E46K α-syn fibril exhibits a novel structure that is distinct from the reported α-syn polymorphs. Compared to the Ac-WT fibril prepared under the same condition, the E46K mutation breaks an important electrostatic interaction of E46-K80 and thus induces the reconfiguration of the overall structure. In the mutant fibril, the FC-N region that used to form the protofilamental interface in the Ac-WT fibril, flips away from the fibril interface and becomes less stable. In line with the structural information, we find that the Ac-E46K fibril is less stable upon both physical and chemical treatments. Furthermore, seeding experiments indicate that the instability and fragmentation-prone property of Ac-E46K fibril may lead to an increased capability of propagation.

Protein structures are determined by their primary sequences. While, regarding amyloid proteins, this dogma becomes more complicated. The reported α-syn fibril structures usually contain the central region of α-syn including residues ~37-99^15,16,22–24,27^. This ~60 a.a. long sequence can fold into various structures to construct polymorphic α-syn fibrils (Fig. 5a). Then, how many fibril structures can α-syn potentially form? What determines the possibilities? We notice that different α-syn structures contain different electrostatic interactions, which play an important role in defining and stabilizing the overall structure (Fig. 5a). In the Ac-WT fibril, intra-molecular electrostatic interactions of K58-E61 and E46-K80 lock the Greek key-like fold of α-syn, and the inter-molecular interactions of E45-H50-E57’ are involved in stabilizing the fibril interface, which pairs α-syn protofilaments to form mature fibrils (Fig. 5a). In contrast, in another WT fibril polymorph^23^, the electrostatic interaction forms between K45 and E57’, which mediates the dimerization of the protofilaments (Fig. 5a). Alternatively, the Ac-E46K fibril contains an intra-molecular electrostatic interaction between K45 and E57, which locks the β-hairpin fold of the FC-N region, and an inter-molecular interaction between E61 and K80’, which stabilizes the interface of the mutant fibril (Fig. 5a). Different patterns of electrostatic interactions are also observed in other amyloid fibrils. For instance, tau fibrils derived from Pick’s disease exhibit different electrostatic interactions from those from AZ and chronic traumatic encephalopathy (CTE)^35–37^ (Supplementary Fig. 7). The structural information suggests that different combinations of charged residues may serve as one of the driving forces for the selection of fibril polymorphs. In a hypothetic five-strand four-charged-residue model, limited topologies can be defined by different combinations of charged residues (Fig. 5b). Thus, although real cases are more complicated, there might be limited and probably predictable variations of stable α-syn folds. In addition, mutations and post-translational modifications can further enrich the potential variations.

**Fig. 5.**
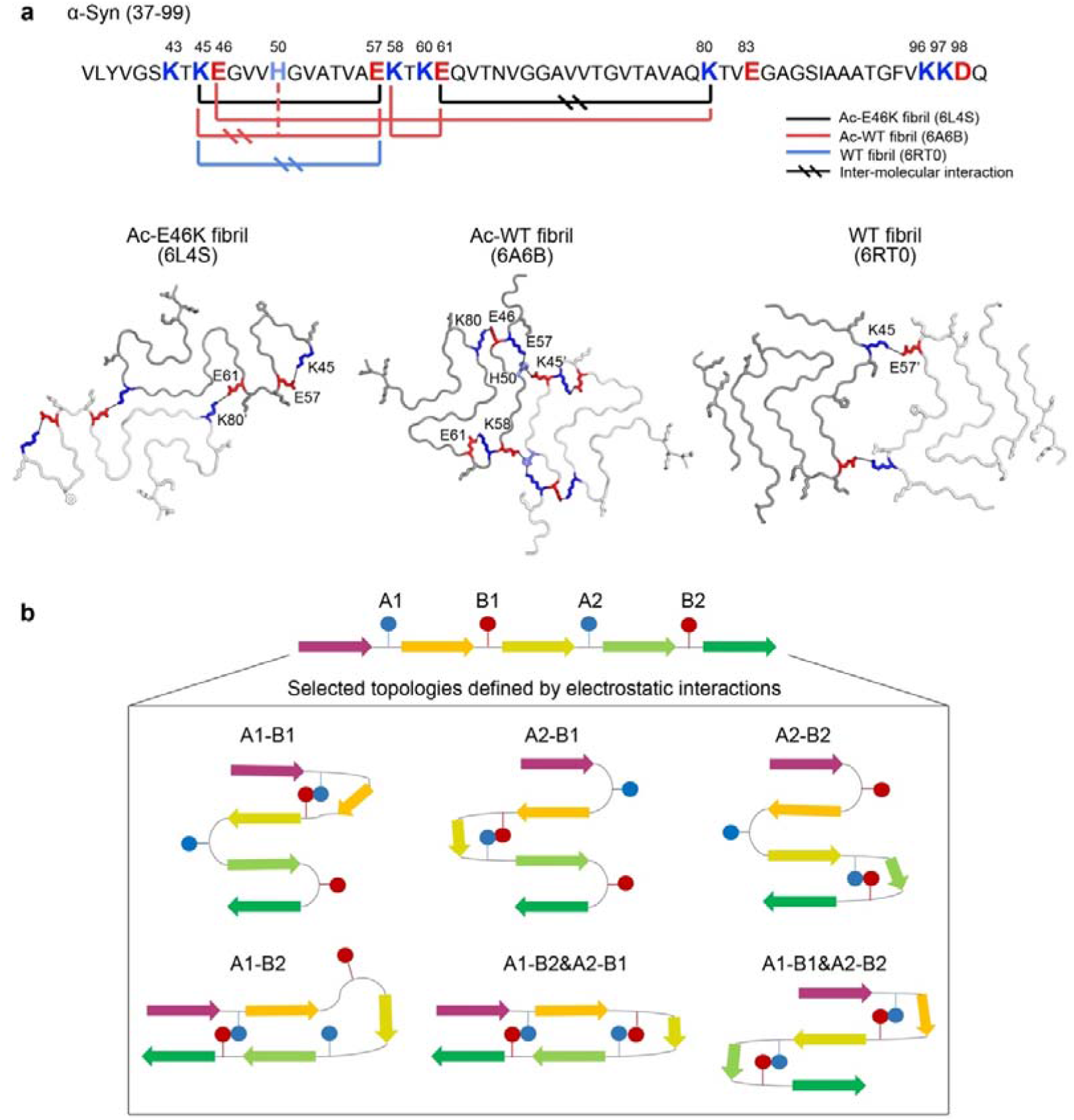
Electrostatic interactions in α-syn fibril polymorphs. **a** Primary sequence of WT α-syn fibril core (FC) is shown on top. K residues are colored in blue; H is light blue; E/D is red. Interacting charged residues in the structures of different α-syn polymorphic fibrils are connected with solid lines. One layer of different polymorphic structures of α-syn fibrils are shown below. The dimeric α-syn molecules are colored in different levels of gray. Electrostatic interactions are highlighted with K colored in blue, H in light blue and E in red. PDB IDs of the structures are provided in parentheses. **b** A hypothetic protein model that contains five β stands, two positively-and two negatively-charged residues (represented by blue and red solid circles, respectively). The positively-charged residues are termed as A1 and A2; the negatively-charged residues are termed as B1 and B2. Combinations of electrostatic interactions include A1-B1, A1-B2, A2-B1, A2-B2, A1-B1& A2-B2, and A1-B2&A2-B1. The schematic demonstrates selected topologies that are defined by these different combinations.

## ACKNOWLEDGEMENTS

This work was supported by the Major State Basic Research Development Program (2016YFA0501902), the Science and Technology Commission of Shanghai Municipality (18JC1420500), “Eastern Scholar” project supported by Shanghai Municipal Education Commission, the Shanghai Pujiang Program (18PJ1404300), and the National Natural Science Foundation (NSF) of China (91853113, 31872716 and 31800709), the Chinese Academy of Sciences, the Advanced Innovation Center for Structural Biology, Tsinghua-Peking Joint Center for Life Sciences. We acknowledge Tsinghua University Branch of China National Center for Protein Sciences Beijing and the cryo-EM platform of Peking University for providing facility supports in cryo-EM.

## AUTHOR CONTRIBUTIONS

C. L., X.L. and D.L. designed the project. K.Z. and H.L. prepared the Ac-WT and Ac-E46K monomer and fibril samples. Z.L., H.L. K.Z. performed the biochemical assays. Z.L. performed the AFM experiments. Y.L and K.Z. performed the cryo-EM experiments. Y.L. built and refined the structure model. All of the authors are involved in analyzing the data and contributed to manuscript discussion and editing. D. L. and C.L. wrote the manuscript.

## Methods

### Preparation of Ac-WT and Ac-E46K α-syn

Protein expression and purification of full length α-syn Ac-WT and Ac-E46K mutation follows the same protocol described previously^15^. Briefly, the WT and E46K α-syn gene were inserted into pET22 vector and transformed into *E. coli* expression cell line BL21(DE3). Fission yeast N-acetyltransferase complex B was co-expressed with α-syn to obtain the N-terminally acetylated α-syn^38^. 1 mM isopropyl-1-thio-D-galactopyranoside (IPTG) was used to induce protein expression at 37°C for 4 h. Cells were harvested in 50nM Tris-HCl, pH 8.0, 1mM phenylmethylsulfonyl fluoride, 1mM EDTA and lysed by sonication. After centrifugation at 16000× g for 30 min, the supernatant was boiled at 100 °C for 10 min. After another centrifugation, streptomycin (20 mg/mL) was added to remove nucleic acid in the supernatant. pH was adjusted to 3.5 using 2M HCl to precipitate other unwanted components. 50mM Tris-HCl (pH 8.0) was used to dialyze the supernatants overnight. Q column (GE Healthcare, 17-5156-01) was used to purify the protein samples and size-exclusion chromatography Superdex 75 (GE Healthcare, 28-9893-33) was used for further purification. After purification, an on-line EASY-nL-LC 1000 coupled with an Orbitrap Q-Exactive HF mass spectrometer was used to verify that the Ac-E46K α-syn protein sample is indeed acetylation (Supplementary Fig. 1).

### Fibrillation of Ac-WT and Ac-E46K α-syn

100 μM recombinant Ac-WT and Ac-E46K α-syn (in 50 mM Tris, pH 7.5, 150 mM KCl) were incubated at 37 °C with constant agitation (900 rpm) in ThermoMixer (Eppendorf) for a week, respectively. To obtain α-syn PFFs, the fibril was sonicated at 20% power for 15 times (1 s/time, 1 s interval) on ice. Then, the α-syn PFFs (0.5%, v/v) was added to 100 μM α-syn monomer, and further incubated at 37 °C with agitation for a week to obtain mature fibrils for cryo-EM and AFM measurement. The mature α-syn fibrils were concentrated by centrifugation (14,000 rpm, 25 °C, 45 min), washed with PBS and sonicated into α-syn PFFs for primary neuron, PK digestion and Circular dichroism experiments. The Ac-WT and Ac-E46K α-syn fibrils was sonicated at 20% power for 4 times to perform the ThT kinetic assay and AFM measurement of the fibril size distribution.

### CD spectroscopy

To detect the stability of fibrils, 20 μM Ac-WT and Ac-E46K α-syn fibrils were incubated at 0□. Fibril samples were picked up at different incubation time for CD measurement. For freeze-thaw assay, fibril samples were incubated in liquid nitrogen with 10 min for fully freezing and dissolved in water at room temperature for 5 min. The secondary structure of the fibril sample was measured by a Chirascan CD spectrometer (Applied Photophysics, UK). Spectra was recorded from 200 to 260 nm at 0□ or room temperature using a cell with a path length of 0.1 cm. All spectra were collected more than 3 times with a background-corrected against buffer blank.

### Atomic force microscopy (AFM)

20 μM Ac-WT and Ac-E46K α-syn fibrils were sonicated for 4 times (1s on, 1s off) on ice, respectively. 8 μl of diluted samples (10 μM) were mounted on a freshly cleaved mica for 3 min and gently rinsed with Milli-Q water for removing unbound fibrils. The dried sample was prepared with nitrogen flow. Images were acquired with ScanAsyst air mode by using Nanoscope V Multimode 8 (Bruker). SNL-10 probe was taken with a constant of 0.35 N/m for scanning. Measurements were recorded at 512*512 pixels at a line rate at 1.5 Hz. The percentage for different length of fibrils were processed and analyzed by using NanoScope Analysis software (version 1.5). As for characterizing the morphology of mature α-syn fibrils, samples were acquired with the concentration of 2 μM. The image and height were analyzed on the Nanoscope software and imageJ.

### Proteolytic digestion of α-syn PFFs

α-syn PFFs (10 μg, in PBS, PH 7.4) were incubated for 30 min at 37 □ in the presence of proteinase K (0.5, 1, 1.5, 2, 2.5 μg/ml, Invitrogen). Reaction solution was terminated by adding 1 mM PMSF. Then the samples were boiled with SDS-loading buffer for 15 min and loaded on 4%-20% Bis-Tris gels (GenScript). The gels were stained by Coomassie brilliant blue and images were recorded and analyzed with Image Lab 3.0 (Bio-Rad).

### Kinetic analysis of the seeded α-syn fibrillization

To evaluate the seeding capability of Ac-WT and Ac-E46K α-syn PFF, we set up a seeded α-syn fibrillization assay by added different concentration of α-syn WT/E46K PFFs (20% power, 4 times of sonication) (0%, 0.3%, 0.5%, mass concentration ratio to α-syn monomer) into 50 μM α-syn monomer (in 50 mM Tris, pH 7.5, 150 mM KCl, 0.05% NaN_3_), respectively. 10 μM Thioflavin-T (ThT) was also added to the reaction mixture. Reactions were performed in triplicate in a 384 well optical plate (Thermo Scientific Nunc). The plate was detected by a Varioskan® Flash microplate reader (Thermo Scientific). Fluorescent intensities of each reaction were acquired by using 440 nm (excitation) and 485 nm (emission) wave-lengths, with a bottom read. Graphing was performed with GraphPad Prism 6.

### Negative-staining electron microscopy

5 μl aliquot of fibril sample was applied to a glow-discharged 200 mesh carbon support film (Beijing Zhongjingkeyi Technology Co., Ltd.) for 45 s. Then the grid was washed with 5 μl double-distilled water and followed by another wash of 5μl 3% w/v uranyl acetate. Another 5 ul 3% w/v uranyl acetate was added to the grid to stain the sample for 45 s. After removing the excess buffer by filter paper, the grid was dried by infrared lamp. The sample imaging was applied by Tecnai T12 microscope (FEI Company) operated at 120 KV.

### Cryo-electron microscopy

An aliquot of 4 μl of 6 μM Ac-E46K α-syn fibril solution was applied to a glow-discharged holey carbon grid (Quantifoil R1.2/1.3, 300 mesh), blotted for 6.5 s, and plunge-frozen in liquid ethane using FEI Vitrobot Mark IV with 95% humidity at 16 °C. The grids were loaded onto a FEI Titan Krios microscope operated at 300 KV with magnification of 105,000×, in which a GIF Quantum energy filter (slit width 20 eV) were equipped. A Gatan K2 Summit camera recorded the cryo-EM micrographs using super-resolution mode with pixel size 0.665 Å. Defocus values were set from −2.4 to −1.5 μm. All of the micrographs were dose-fractionated to 32 frames and the electron dose rate was set to ~8.2 counts/physical pixel/s (~6.25 e^-^/s/Å^2^), total exposure time is 8 s, and 0.25 s per frame. Hence, the total dose is ~50 e^-^/Å^2^. AutoEMation2 written by Dr. Lei was used for all data collection^39^.

### Image pre-processing

All 32 frames were aligned, summed and dose-weighted by MotionCorr2 and further binned to a pixel size of 1.33 Å^40^. The defocus values of dose-weighted micrographs were estimated by CTFFIND4^41^, and micrographs which maximum resolution at where the Thon ring with good fit up to is higher than 6 Å were kept for the subsequent processing in RELION3.0^42^. 13064 filaments were picked manually from 754 micrographs. The exacted segments (box size: 160×160, inter-box distance of 19 Å) were classified by 2D (two-dimensional) classification. Segments of classes that have clear cross-beta strands kept, and a steric zipper conformation exists which is same with the conformation of the wild type of alpha-synuclein fibril.

### Helical parameters determination

Twist angle and rise value of helical polymer was calculated before reconstruction. Micrographs, which were involved in the selected 2D classes, were used to estimate the rise value by a C++ program that searches the strongest diffraction position around the 1/4.8 Å^-1^ in Fourier space. The average of these positions was converted to the rise value in real space. The way to estimate the twist angle was the same as that described in our previous structural determination of Ac-WT α-syn fibril^15^, in which helical pitch length of each filament was estimated and the average of these helical pitch lengths was used to calculate the twist angle.

### Helical reconstruction

Several iterations of 2D classification, 3D classification and 3D refinement were performed to filter the segments belonging to the same conformation. Segments of the selected 2D classes were used to reconstruct a reference with more feature in 3D refinement from the initial featureless cylinder model created by the relion_helix_toolbox program. 3D classification (k=2) then built a reasonable map based on the improved reference model. 3D refinement of selected 3D class improved the information map with higher resolution. The final map was convergence with the rise of 2.376 Å and the twist angle of −179.371°. The map was sharpened with a B-factor of −97.8051 Å^2^ using post-processing subroutine in RELION. The overall resolution was reported at 3.37 Å by the gold-standard FSC = 0.143 criteria. Local resolution estimation calculated by ResMap^43^.

### Polymorphs of Ac-E46K α-syn fibril

Two polymorphs at least existed in micrographs. One is twisted filament, and the another one is non-twisted filament which currently cannot be solved since this polymorph has no twist. All filament including twisted and non-twisted filaments in 754 micrographs were count as 18891. Among them, 13064 was twisted filaments which was used for reconstruction. The percentage of twisted filament is 69.15% and was dominated in the two fibril polymorphs.

### Model building and refinement

A homology model based on the NMR structure (PDB entry code 2N0A) was built and modified by COOT^28,44^. The model with 3 adjacent layers (6 promoters) was refined using the real-space refinement program in PHENIX^45^. The subunit dimers in the middle of 3 layers was extracted and used as the final model.

### Accession codes

Density maps of the mAc-E46K α-syn fibril are available through EMDB with entry code: EMD-0833. The structural model was deposited in the Protein Data Bank with entry code: 6L4S. The data that support the findings of this study are available from the corresponding authors upon request.

