## Supplementary material for "Parkinson’s disease associated mutation E46K of α-synuclein triggers the formation of a novel fibril structure"

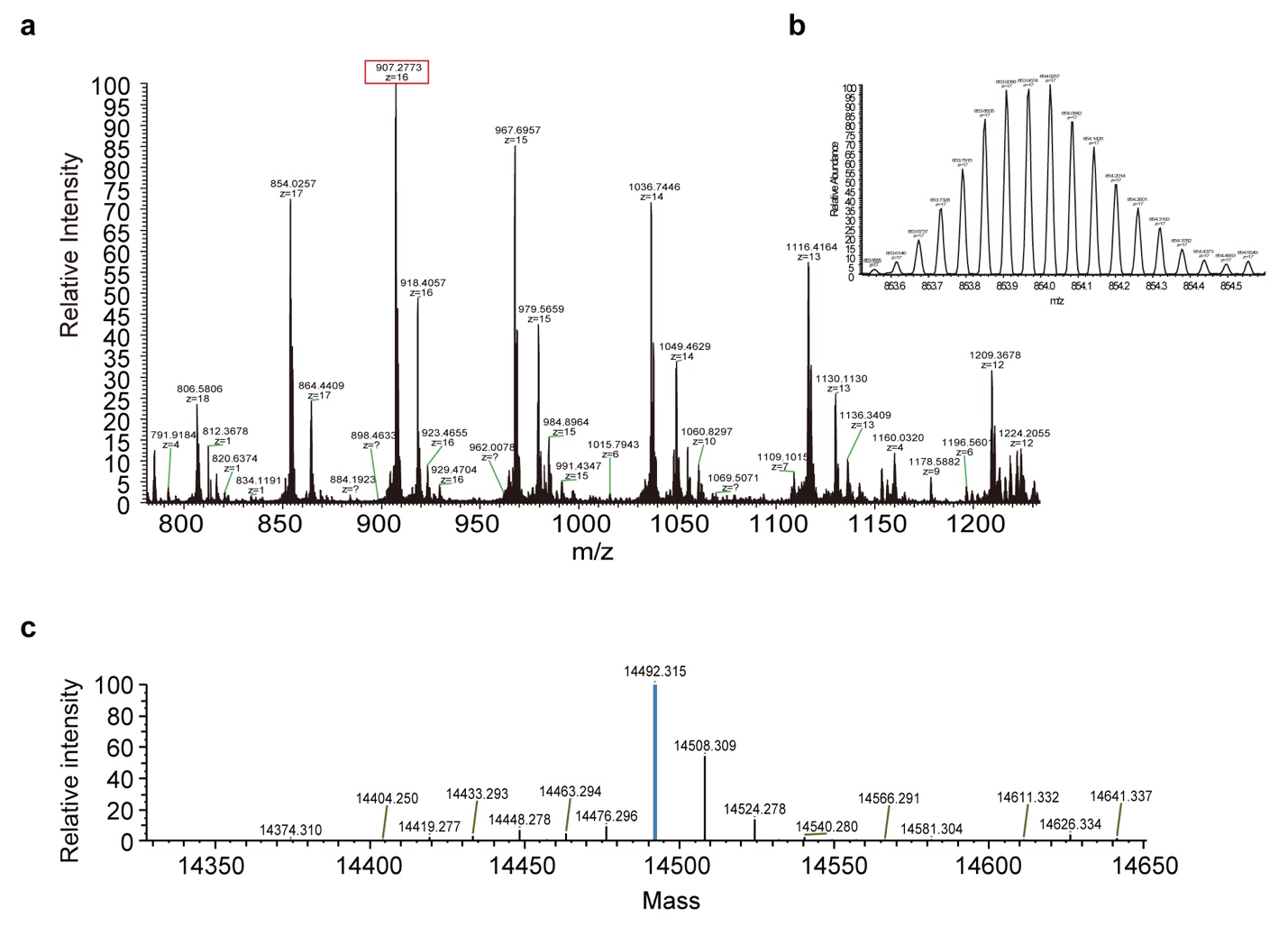

**Supplementary Figure 1** MS validation of the recombinant Ac-E46K α-syn. **a** The different charge states of the Ac-E46K α-syn. **b** The isotopically resolved profile of the Ac-E46K α-syn. **c** The deconvoluted spectrum confirms that the protein is acetylated.

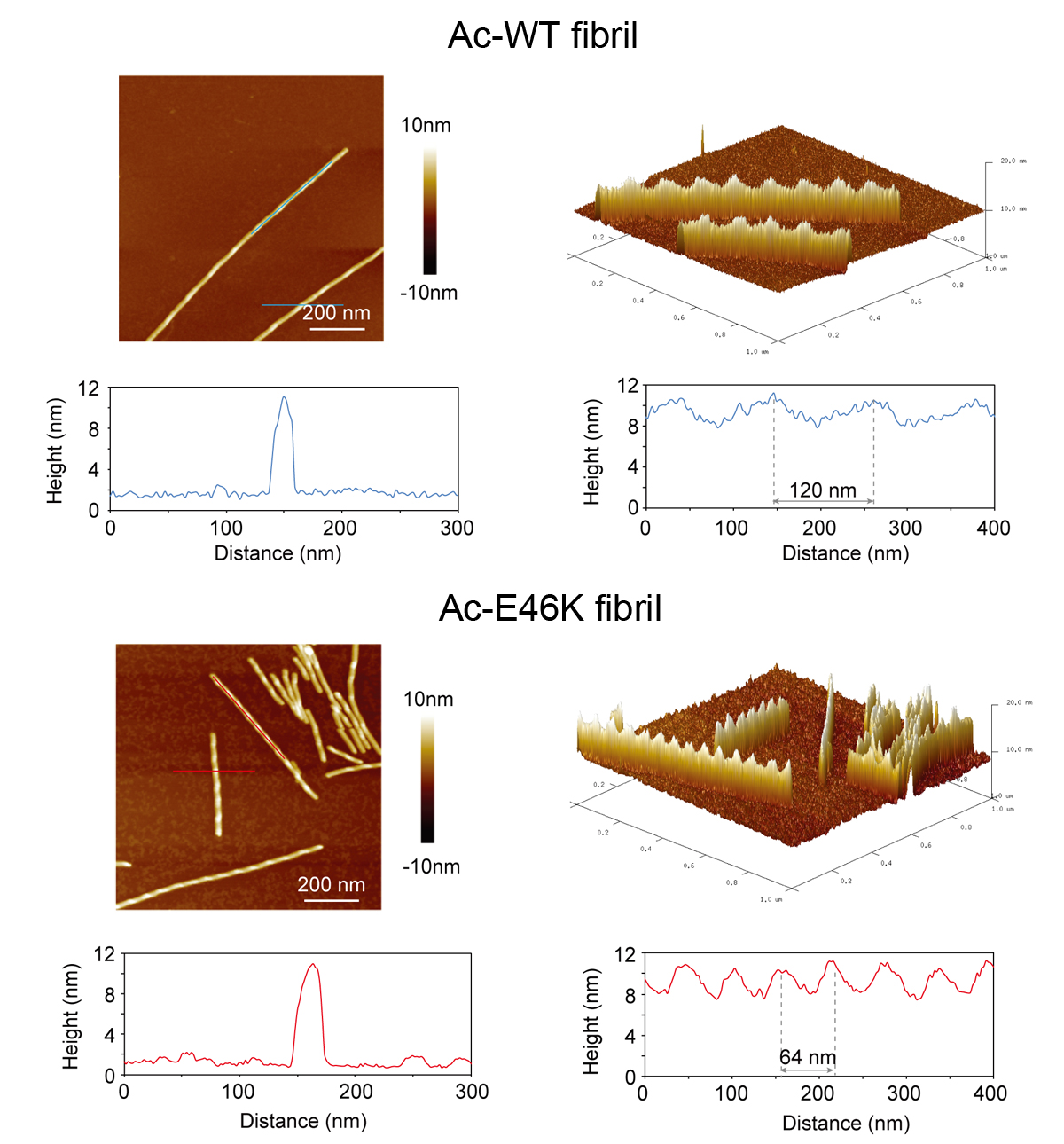

**Supplementary Figure 2** AFM measurement of the Ac-WT and Ac-E46K α-syn fibrils. The fibrils were formed under the same condition: 50 mM Tris, 150 mM KCl, pH 7.5 at 37°C with agitation. AFM 2D and 3D images are shown. Analyses of the cross section and along the fibril are indicated with blue/red lines, respectively. The width and twist periodicity of the fibrils are measured.

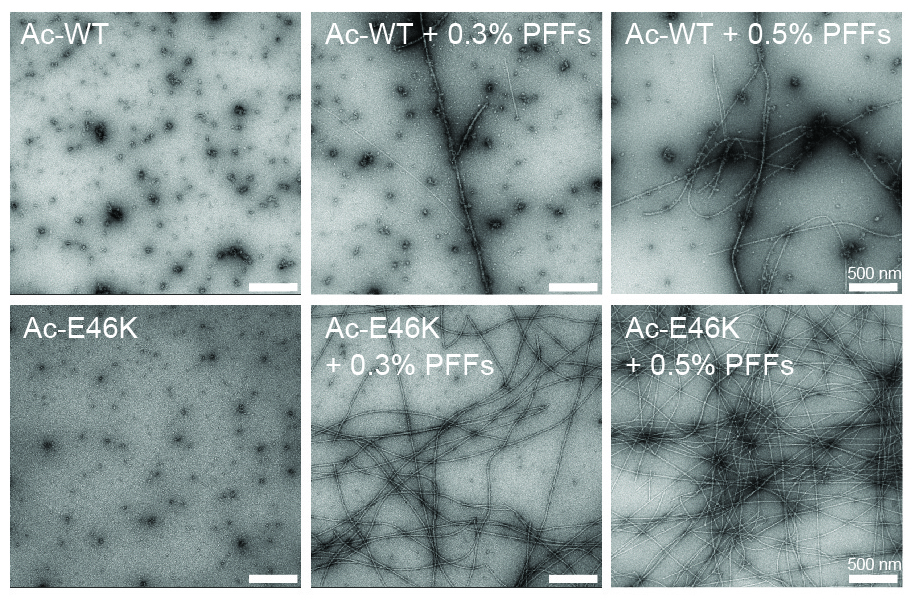

**Supplementary Figure 3** Negative-staining TEM images of Ac-WT and Ac-E46K fibrils seeded with sonicated preformed fibrils (PFFs). Images were taken at 80 h. Scale bar: 500 nm.

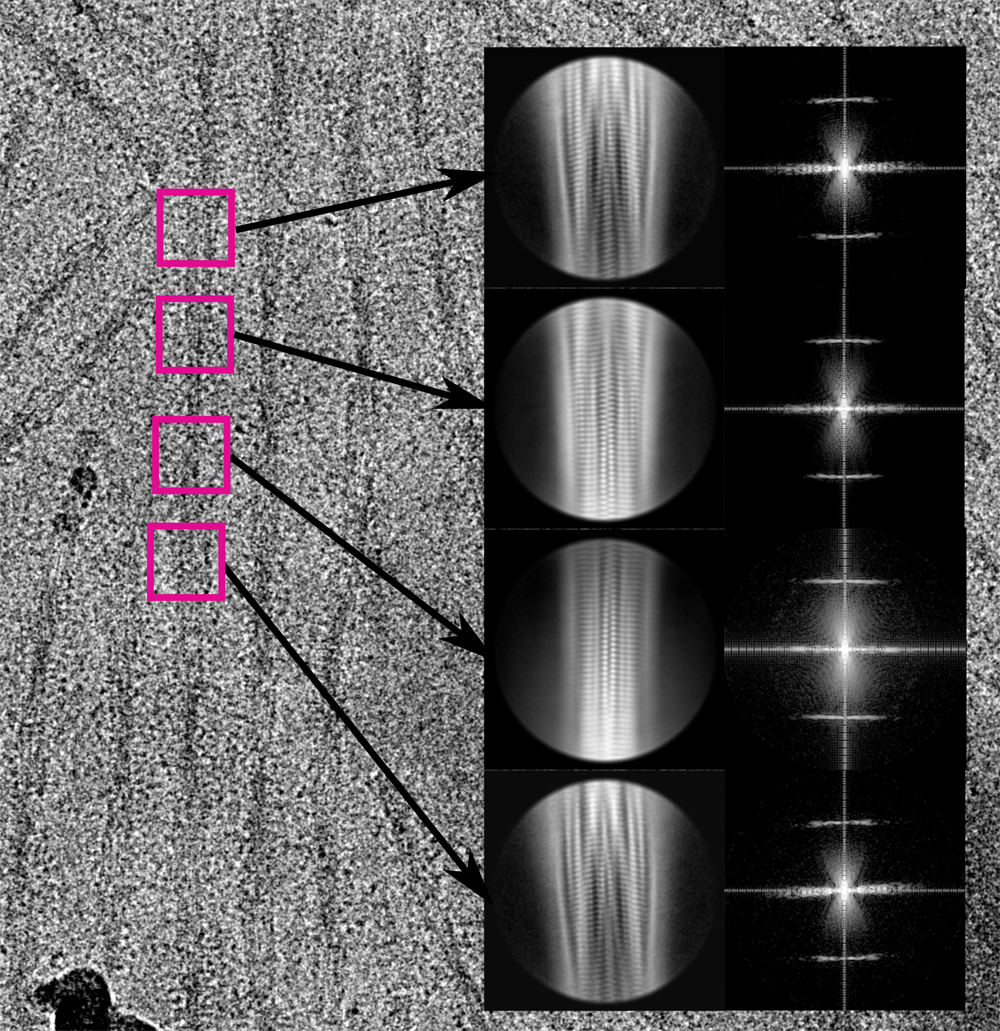

**Supplementary Figure 4** 2D classification of the Ac-E46K fibril. A cryo-EM micrograph of the Ac-E46K fibril is shown. The zoom-in views show the 2D class averages and power spectra of the indicated fibril segments.

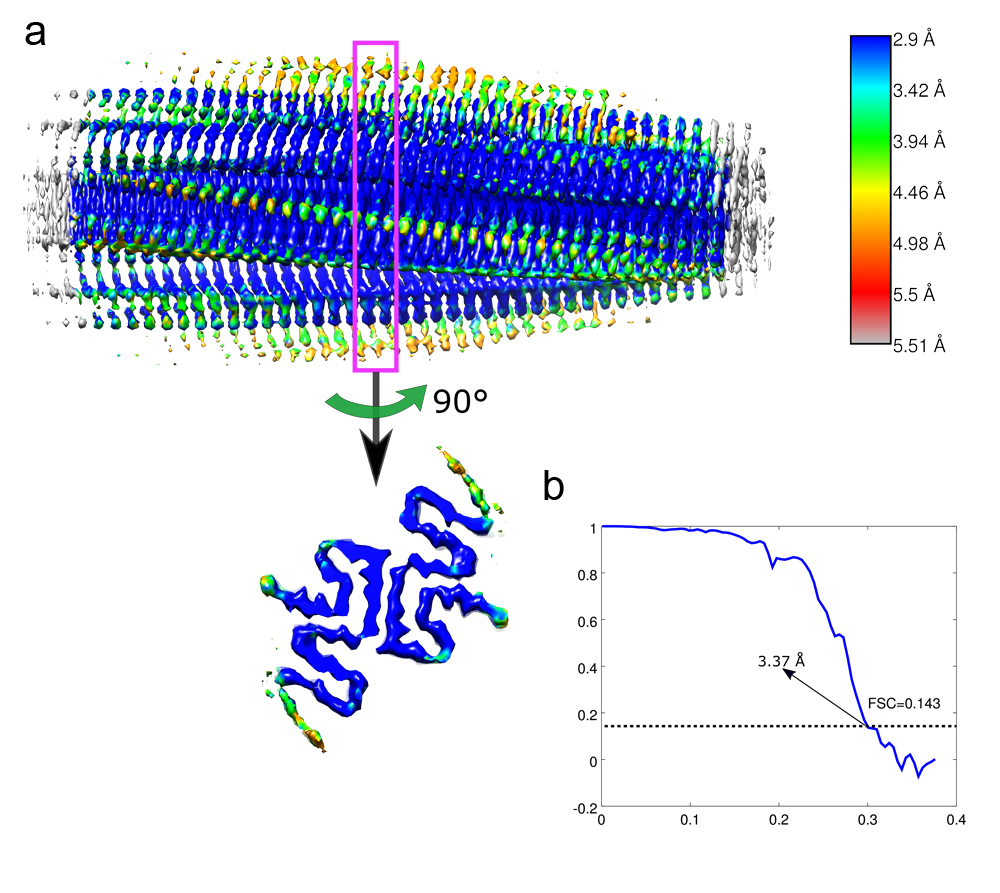

**Supplementary Figure 5** Resolution estimation of the cryo-EM structure of Ac-E46K α-syn fibril. **a** Local resolution estimation. EM reconstruction maps are colored based on the local resolutions. The color scale indicates the resolution ranging from 2.90 Å to 5.51 Å. **b** Gold-standard Fourier shell correlation curve of the Ac-E46K α-syn fibril. The overall resolution is 3.37 Å.

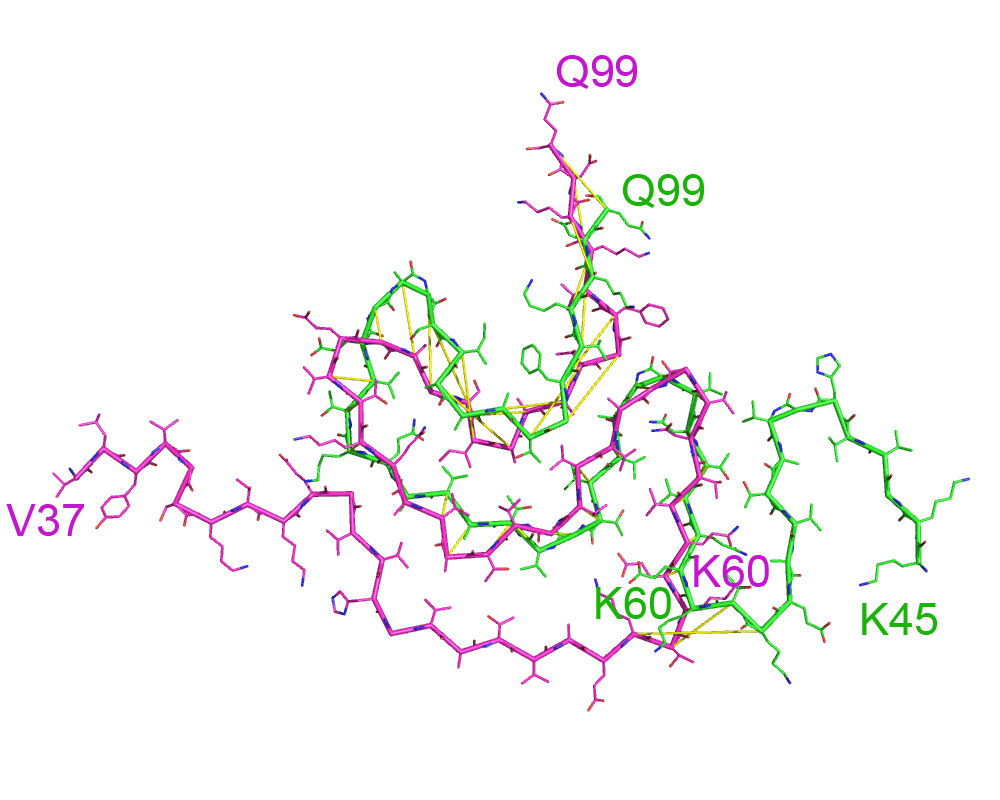

**Supplementary Figure 6** Alignment of the Ac-E46K (green) and Ac-WT (magenta) α-syn subunit structures. The PDB ID of the Ac-WT fibril structure is 6A6B. The FC-N regions of the two structures diverge at K60. The FC-C regions of the two structures adopt a similar topology, while their structural similarity is low with an r.m.s.d. of Cα atoms of 5.558 Å. Yellow lines highlight the discrepancy between the same residues of the two structures.

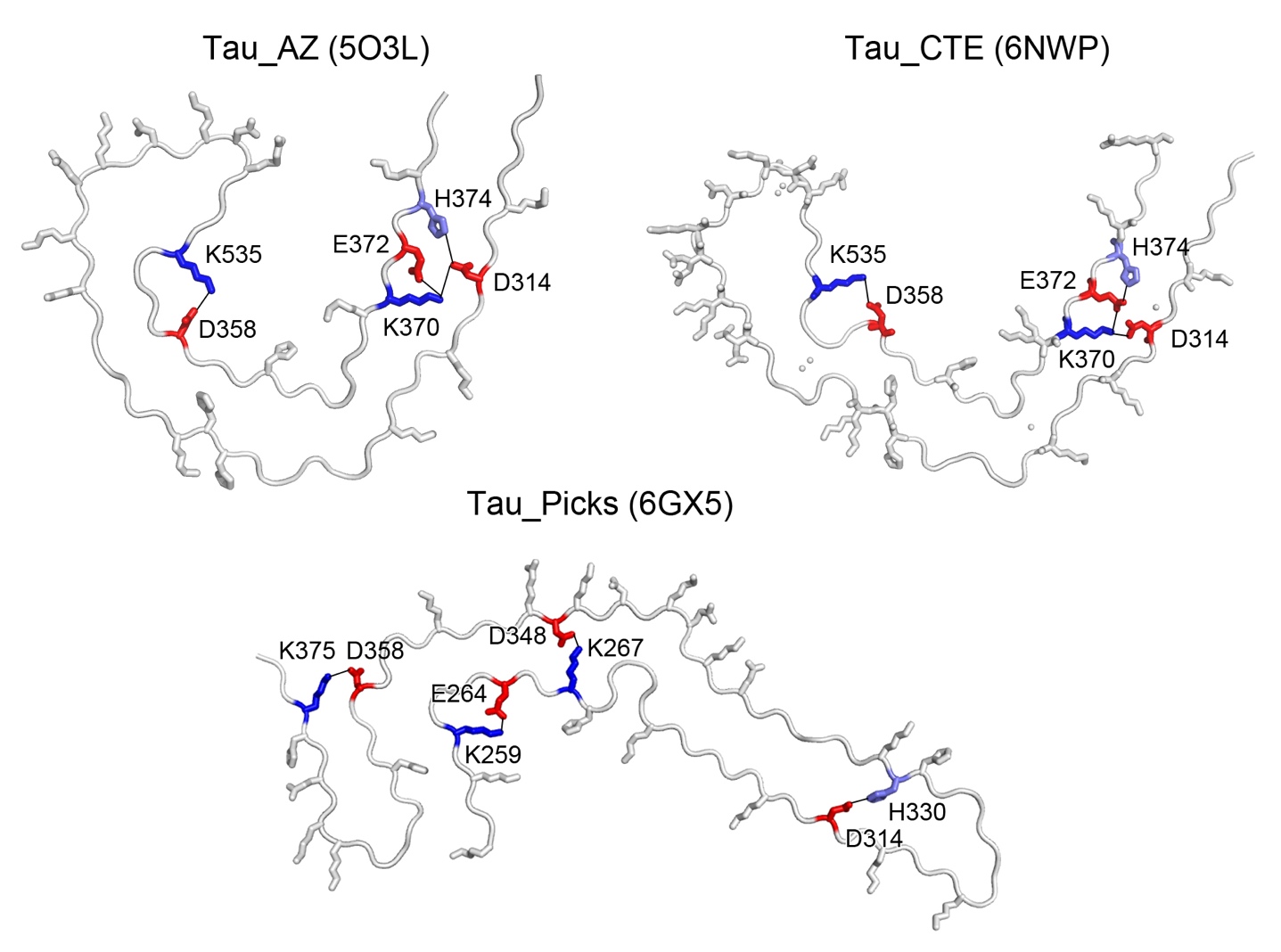

**Supplementary Figure 7** Electrostatic interactions in tau fibril polymorphs. One layer of different polymorphic structures of tau fibrils. Single tau subunits are shown since there is no inter-molecular electrostatic interaction in the tau fibrils reported so far. Electrostatic interactions are highlighted with K colored in blue, H in light blue and E/D in red. PDB IDs of the structures are provided in parentheses.
